# Selection bias in instrumental variable analyses

**DOI:** 10.1101/192237

**Authors:** Rachael A. Hughes, Neil M. Davies, George Davey Smith, Kate Tilling

**Affiliations:** Population Health Sciences, Bristol Medical School, University of Bristol, Bristol, UK; MRC Integrative Epidemiology Unit, University of Bristol, Bristol, UK

**Keywords:** causal exposure effect, collider stratification bias, instrumental variable, selection bias, two stage least squares

## Abstract

Participants in epidemiological and genetic studies are rarely true random samples of the populations they are intended to represent, and both known and unknown factors can influence participation in a study (known as selection into a study). The circumstances in which selection causes bias in an instrumental variable (IV) analysis are not widely understood by practitioners of IV analyses. We use directed acyclic graphs (DAGs) to depict assumptions about the selection mechanism (factors affecting selection) and show how DAGs can be used to determine when a two-stage least squares (2SLS) IV analysis is biased by different selection mechanisms. Via simulations, we show that selection can result in a biased IV estimate with substantial confidence interval undercoverage, and the level of bias can differ between instrument strengths, a linear and nonlinear exposure-instrument association, and a causal and noncausal exposure effect. We present an application from the UK Biobank study, which is known to be a selected sample of the general population. Of interest was the causal effect of education on the decision to smoke. The 2SLS exposure estimates were very different between the IV analysis ignoring selection and the IV analysis which adjusted for selection (e.g., 1.8 [95% confidence interval *−*1.5, 5.0] and *−*4.5 [*−*6.6*, −*2.4], respectively). We conclude that selection bias can have a major effect on an IV analysis and that statistical methods for estimating causal effects using data from nonrandom samples are needed.

## Introduction

The main aim of many epidemiological studies is to estimate the causal effect of an exposure on an outcome. Instrumental variable (IV) analyses are increasingly used to overcome bias due to unmeasured confounding. An IV analysis requires a variable, known as the instrument, to satisfy three assumptions: the instrument is associated with the exposure, the instrument only causes the outcome to change via its impact on the exposure, and there is no confounding between the instrument and the outcome [1, 2, 3]. Based on the observed data, the first IV assumption can be tested, but the latter two are untestable [4].

As with any statistical analysis, inference about the causal exposure effect (here onwards shortened to exposure effect) may be invalid when the sample included in the analysis is not a representative (i.e., random) sample of the target population. This could be due to: selection into the study, participant dropout, loss to follow-up, subgroup analysis, or missing data (i.e., restricting the analysis to those participants with complete data). In all of these cases there may be both known and unknown factors that influence the “selection” of participants for analysis.

Following Hernán and Robins [5], we consider selection bias to be distinct from confounding. Confounding is due to the presence of common causes of the outcome and exposure. In contrast, selection bias is due to conditioning on common effects of the outcome and exposure, and is a type of collider-stratification bias [6, 7]. The IV estimate of the causal exposure effect in the study sample is biased by selection when it systematically differs to the value of the exposure effect in the target population [8]. Selection bias is concerned with the internal validity of a study, as opposed to external validity (using a study’s results to make inferences about populations that differ from the target population) [9, 10, 11]. Internal validity is essential before external validity can be considered.

Although selection bias is understood in the methodological literature (e.g., [6, 12, 5]), it is seldom acknowledged in IV analyses or discussed in guidelines for IV analysis (e.g., [13, 14, 15, 16, 17]). However, recent exceptions include examples where selection depends on the: exposure plus confounder, or outcome [18], exposure [19, 18], instrument plus measured and unmeasured confounders (of the outcome-exposure association) [20], exposure and measured variable (which causes the outcome) [21], missing values of measured covariates [22, 23], and unmeasured confounder plus measured covariates [24].

In the IV literature a small number of papers have used directed acyclic graphs (DAGs) [25, 26, 27] to illustrate when selection violates the assumptions of an IV analysis [19, 20, 21, 18]. However, these papers cover a limited range of selection scenarios, with Gkatzionis and Burgess [18] confining their discussion to Mendelian randomisation, and Ertefaie et al, and Canan et al [20, 21] provide an incomplete explanation of the consequence of selection. Only one paper [18] considered if the effects of selection differed according to a null and non-null causal exposure effect, and none of these papers investigated if the consequences of selection differed according to a linear and nonlinear exposure-instrument association.

We use DAGS to illustrate the circumstances in which an IV analysis is biased by selection for a wide range of selection scenarios which can occur in practice. Via simulations, we show how the consequences of selection can depend on the factors determining selection, strength of the instrument, whether the causal effect is null or not null, and linearity of the exposure-instrument association. Finally, using a real application we show how an IV analysis ignoring nonrandom selection can reach different to an IV analysis which adjusts for nonrandom selection.

## When does selection lead to bias?

### Description of our IV analysis

We want to estimate the effect of a continuous exposure *X* on a continuous outcome *Y*, and we denote this causal exposure effect by *β_X_*. The *Y* – *X* association is confounded by unmeasured variables *U* and measured variables *C*. In the full sample (i.e., the selected and unselected participants), the instrument *Z* satisfies the three assumptions of an IV analysis (without conditioning on *C*).

To identify *β_X_* we assume homogeneous exposure effects (i.e., *β_X_* is the same for all individuals [15]. We estimate *β_X_* using the two stage least squares (2SLS) method [28] and denote its 2SLS estimate by 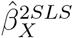. In the first stage of 2SLS, *X* is regressed on *Z* to give fitted values *X̂.* In the second stage, the regression coefficient of *Y* on fitted values *X̂* is the 2SLS estimate, 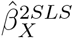. When *Z* is a single instrument, 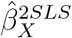 is equivalently estimated using the ratio of coefficients method [29, 30].

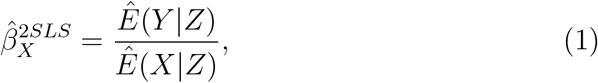

where the numerator, *Ē*(*Y* | *Z*), is the estimated coefficient from the regression of *Y* on *Z*, and the denominator, *Ē* (*X* | *Z*), is the estimated coefficient from the regression of *X* on *Z*. We also estimate the exposure effect conditional on measured confounders *C*, and denote this conditional 2SLS estimate by 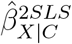.

### Selection mechanisms

Whether 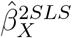 is biased by selection depends on the reasons for selection, i.e. the “selection mechanism”. Figures 1a to 1i depict DAGs showing the causal relationships among the variables of our IV analysis under nine selection mechanisms, where *S* is a binary variable indicating whether a participant is selected or unselected. Restricting the analysis to the selected sample implies conditioning on *S* which is represented by a box around *S*. Because a DAG is non-parametric, the discussion below is not specific to continuous variables only; for example, they also apply when *X*, *Z* and *Y* are binary variables and examined on the risk difference scale (as in our applied example below). Unless otherwise stated, whether 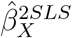 is biased by selection equally applies when the true causal effect is null and not null. Also, in our example all variables are measured without error - however, whether 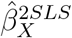 is biased by selection equally applies when selection depends on variables measured with error [5].

Table 1 summarises when 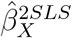 and 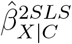 are biased by selection. When selection is completely at random (Figure 1a), or depends on *Z* (Figure 1b), or *U* (Figure 1c), 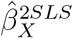 and 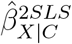 are not biased by selection. Here, selection does not imply conditioning on a collider (nor a descendant of a collider), and so the IV assumptions remain true in the selected sample.

**Figure 1:**
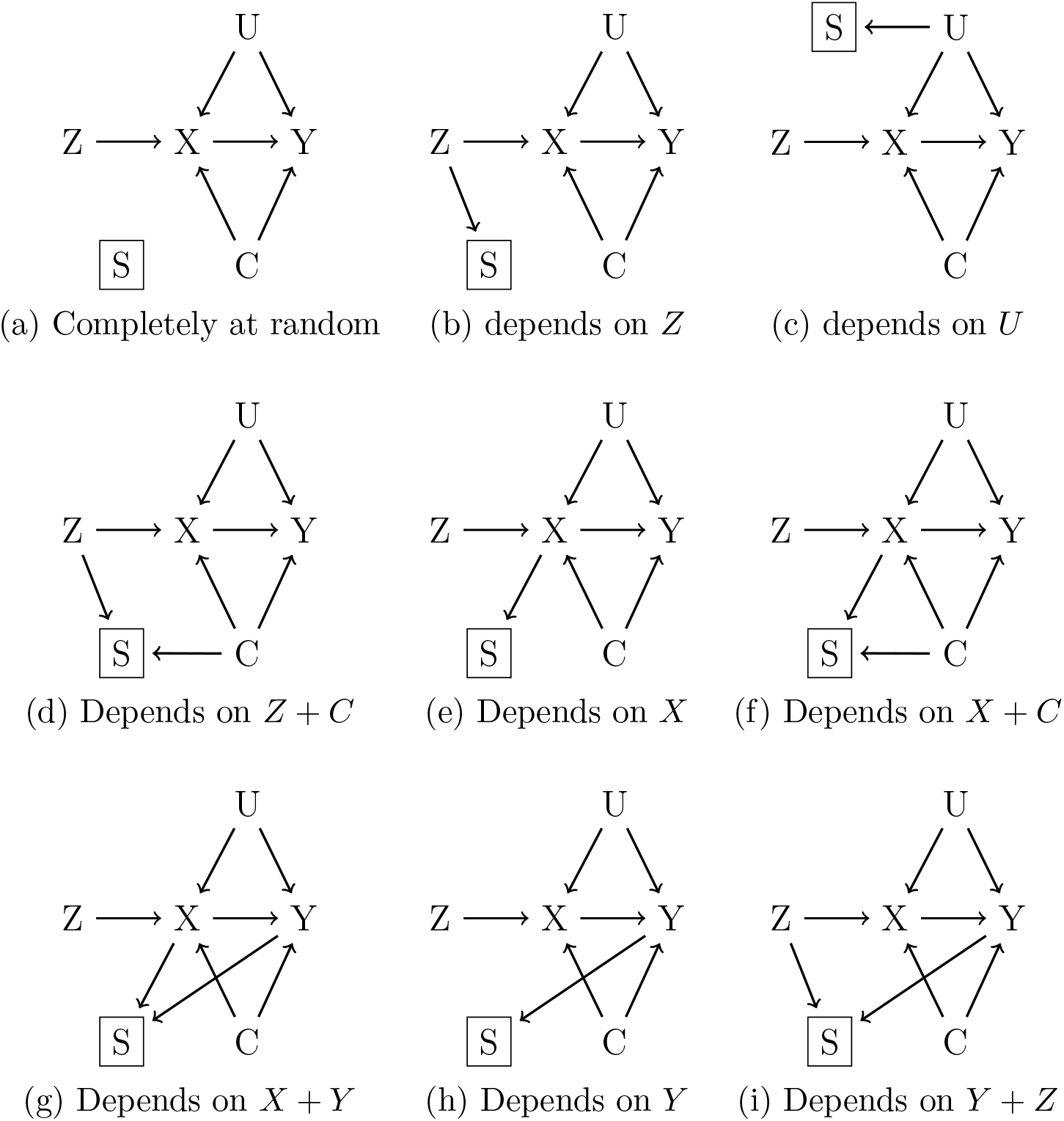
Directed acyclic graphs of an instrumental variable analysis under nine different selection mechanisms.

**Table 1.**
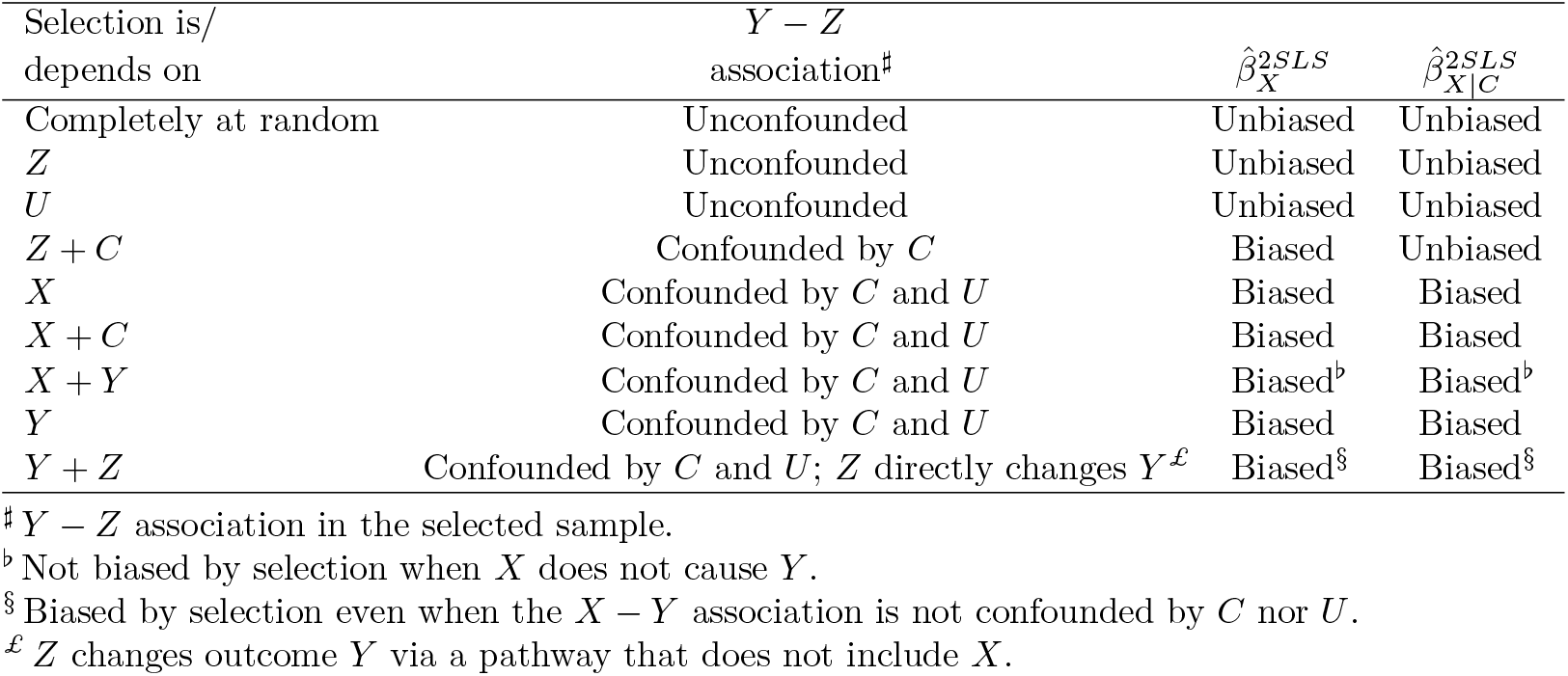
Potential bias of the two stage least squares (2SLS) estimate of the casual exposure effect, 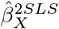, and the corresponding 2SLS estimate conditional on *C*, 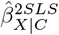, according to different selection mechanisms.

When selection depends on *Z* + *C*, *X*, *X* + *C*, *X* + *Y* or *Y* (Figures 1e, 1f, 1g or 1h, respectively), 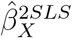 is biased by selection because the *Y* − *Z* association becomes confounded in the selected sample. Here, selection implies conditioning on a collider which opens a noncausal pathway between *Z* and *Y* via a confounder (e.g., selection on *X* + *C* opens pathway 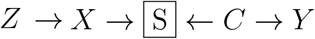). For selection mechanism *Z* + *C*, the *Y* − *Z* association is confounded by *C* only. Therefore, whilst 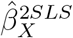 is biased by selection on *Z* +*C*, 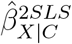 is not biased because the only noncausal pathway is via *C* which is re-blocked by conditioning on *C*. For the other selection mechanisms, the *Y* – *Z* association is confounded by *C* and *U*. Therefore, whilst estimating 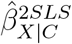 reduces the level of selection bias (by eliminating confounding by *C*), 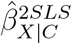 remains biased because the *Y* – *Z* association is still confounded by *U* in the selected sample.

Selection depending on *Y* has the special property that 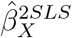 and 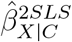 are only biased by selection when *X* causes *Y* (the true exposure effect is not null). When *X* does not cause *Y*, the pathways between *Z* and *Y* via *C* and *U* are blocked by the absence of an edge between *X* and *Y* (e.g., Z → *X Y* ← *U*).

When selection depends on *Y* +*Z* (Figure 1i), 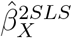 and 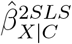 are biased by selection because the instrument is directly associated with the outcome (i.e., there exists a *Y* – *Z* association which is not via *X*) in the selected sample. Here, selection implies conditioning on collider *S* which unblocks pathway 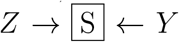. When *X* causes *Y*, selection depending on *Y* + *Z* also results in violating a second IV assumption because the *Y* – *Z* association is confounded by *C* and *U* in the selected sample (as discussed for selection on *Y* only).

A more detailed discussion is given in the appendix.

## Simulation study

For our IV analysis example, we investigated the effects of different selection mechanisms on 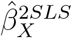. We excluded selection on *U* because it is similar to selection on *Z*, and excluded selection on *Y* + *Z* because we considered it less likely to occur in practice.

### Methods

We simulated data on *X*, *Y*, *Z*, *C* and *U* under a multivariate normal distribution, ensuring the three IV assumptions held true in the full sample. Selection was imposed using a logistic regression model, where the covariates of the model included one or more of *X*, *Y*, *Z* and *C* (depending on the selection mechanism). For all selection mechanisms, close to 60% of the participants were selected. We used Stata (StataCorp; Texas, USA) command *ivregress* to perform 2SLS estimation.

We repeated the simulation study for: a causal exposure effect of 1, and a noncausal exposure effect (i.e., 0). A strong instrument (partial 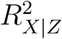 to 0.39 in the full sample), and a moderate instrument (partial 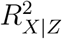 close close to 0.045 in the full sample). A linear *X* – *Z* association (*X* as a function of *Z*) and a nonlinear *X* – *Z* association (*X* as a function of *Z* and *Z*^3^). For all combinations of the simulation settings we generated 3, 000 simulated datasets, each with 20, 000 participants for the full sample.

Of interest was the bias of 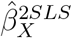, the relative error of its standard error compared to the empirical standard deviation of 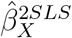, and the coverage of the 95% confidence interval (CI) for 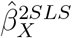. Similarly, for the conditional estimate, 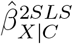. Evidence of systematic bias (i.e., estimates systematically differ from the true value) occurs when the Monte Carlo 95% CI for the bias (bias ± 1.96 × Monte Carlo standard error) excludes zero. Also, based on 3000 simulations the Monte Carlo standard error for the true coverage percentage of 95 is 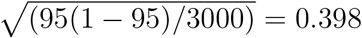 [31], implying that the estimated coverage percentage should lie within the range of 94.2 and 95.8 (with 95% probability). We analysed the simulation results using the *simsum* command [32].

## Results

When there was no selection (the full sample), 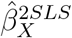 was unbiased and CI coverage was nominal (close to 95%) in all cases, as expected (see Appendix Tables 3 to 6). Figure 2 show the bias of the 2SLS estimates (shown by scatter points; right y-axis) and CI coverage (shown by bars; left y-axis) according to the 8 selection mechanisms and instrument strengths moderate and strong, when the true exposure effect was 1: Figure 2a corresponds to linear *X* – *Z*, and Figure 2b to nonlinear *X* – *Z*. Full results are reported in the corresponding Appendix Tables 3 and 4, respectively.

**Figure 2:**
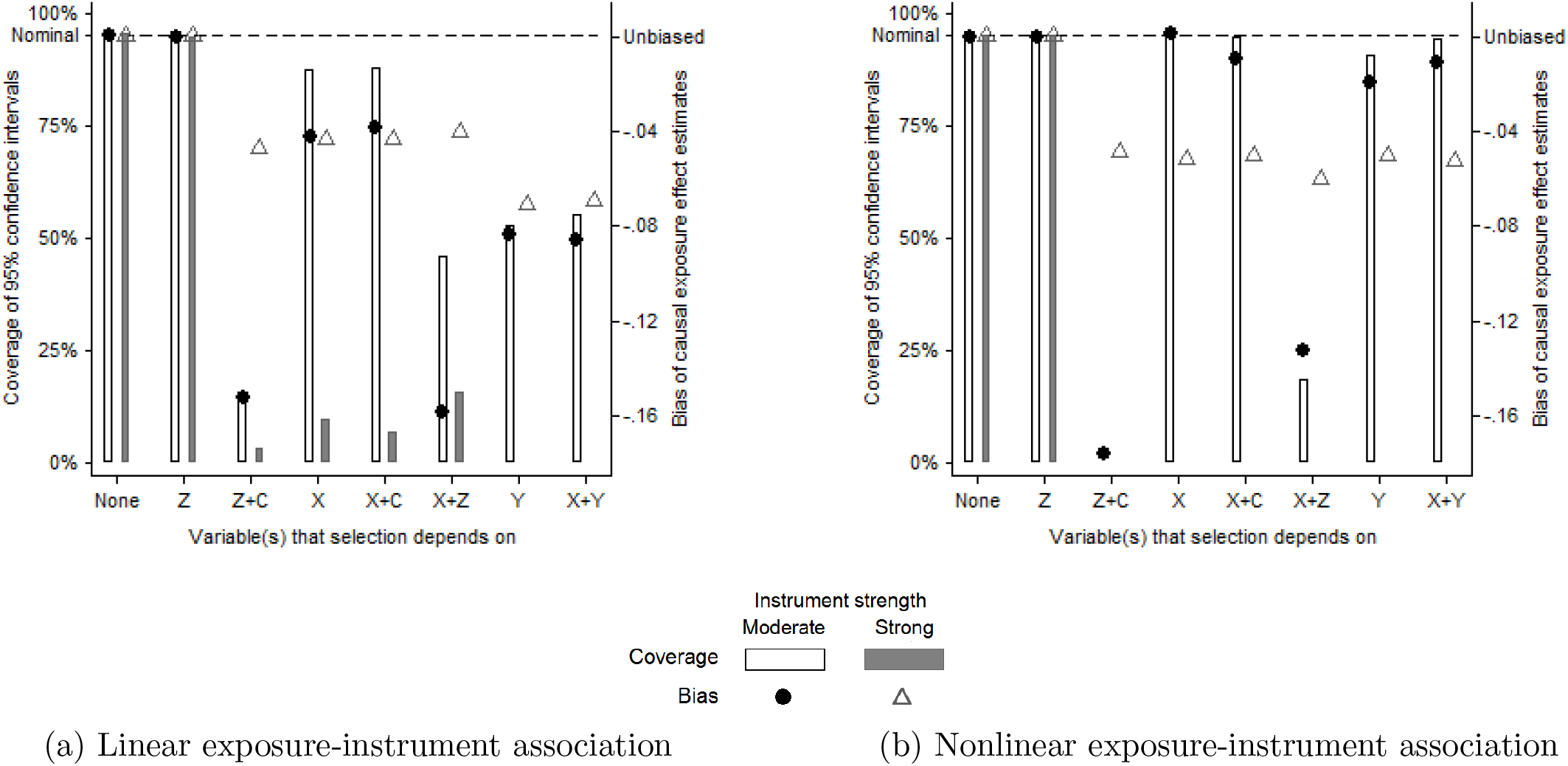
Bias of the two stage least squares estimates (scatter points; right y-axis), and coverage of their 95% confidence intervals (bars; left y-axis) according to different selection mechanisms, and instrument strengths moderate and strong. Panels (*a*) and (*b*) correspond to linear and nonlinear exposure-instrument association, respectively. The true value of the causal exposure effect was 1.

When selection was completely at random (represented as “none”) or depended on *Z* only, 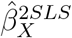 was unbiased and CI coverage was nominal. Because this finding applied to all simulation settings we shall not discuss these two selection mechanisms further. For the remaining selection mechanisms, 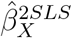 was negatively biased with poor (88%) to severe (0%) CI undercoverage (shown by the absence of a bar) for linear *X* − *Z* (figure 2a).

As expected, selection depending on *Y* did not bias 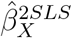 when the exposure effect was null, both for linear and nonlinear *X* −*Z* (Appendix Tables 5 and 6). For the remaining selection mechanisms, the results for a causal and noncausal exposure effect were very similar.

### The impact of instrument strength

When selection partly depended on *Z* (selection mechanisms *Z* + *C* and *X* +*Z*) the level of bias increased with decreasing instrument strength. When selection did not depend on *Z* there were only small differences in the level of bias between the instrument strengths. However, for all selection mechanisms, decreasing the instrument strength resulted in higher CI coverage due to larger standard errors.

### Nonlinear versus linear *X* − *Z* association

In general, the results for nonlinear *X* – *Z* (Figure 2b) follow the same patterns noted for linear *X* – *Z*. Differences in the level of bias between linear and nonlinear *X* – *Z* were far larger for the moderate instrument than the strong instrument because (due to the design of the simulation study) the strength of the nonlinearity was the same for the moderate and strong instruments.

For selection mechanism *Z* + *C*, the effect of the nonlinearity was to decrease the instrument strength, thus increasing the level of bias: when the instrument was moderate the level of bias was 15% higher and the instrument strength (i.e., partial 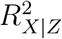) was 17% lower for nonlinear *X* − *Z* compared to linear *X* – *Z* (Appendix table 4). Conversely, for selection mechanism *X*, when the instrument was moderate, the level of bias was 36 times smaller for nonlinear *X* – *Z* compared to linear *X* – *Z*. Nonlinearity caused a large change in the distribution of *X* among the selected participants, and this change in the distribution of *X* resulted in weakening the induced *Z* – *C* and *Z* – *U* associations (i.e., magnitudes close to zero); hence, the large reduction in bias. A similar pattern was noted for the other selection mechanisms where bias resulted from conditioning on collider *X* (or a descendant of *X*).

For the moderate and strong instruments, the standard errors of 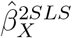 were smaller for nonlinear *X* – *Z* than linear *X* – *Z*, with larger differences for the strong instrument. Consequently, when the level of bias was comparable between linear and nonlinear *X − Z*, CI coverages were poorer for nonlinear *X* − *Z* due to the smaller standard errors. However, in situations where a non-linear *X* −*Z* lowered the level of bias (e.g., selection on *X*) then CI coverages were substantially higher for nonlinear *X* − *Z* despite smaller standard errors.

### The exposure effect conditional on *C*

As expected, for selection mechanism *Z* + *C* 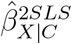 was unbiased and CI coverage was nominal for all simulation settings (Appendix Tables 8 to 11). For the remaining mechanisms, the level of bias for 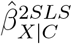 was between 1.2 to 4.5 times lower than that of 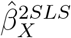, and CI coverage for 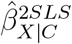 was up to 3 times higher. Otherwise, the results for 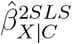 follow the same patterns noted 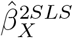.

## Applied example

We conducted an IV analysis to ascertain whether leaving school before age 16 had a causal effect on the decision to smoke [**?**] using data from the UK Biobank study [33], where there is evidence of non-random selection [34]. See the appendix for a detailed description of the analysis.

The binary outcome *Y* was equal to one for ever smokers (included ex-smokers and current smokers), and equal to zero for never smokers. We also considered a second binary outcome, equal to one for current smokers, and equal to zero for ex-smokers and never smokers. Separate analyses were performed on each outcome using the same exposure and instrument. The binary exposure *X* was equal to one if the participant had left school age 16 or older, and equal to zero otherwise. We used a policy reform (often referred to as ROSLA, Raising of School Leaving Age) as an instrument for time spent in education. The binary instrument *Z* was equal to one if the participant turned 15 after the policy reform was introduced, and equal to zero otherwise. There were some measured confounders, *C*, of the exposure-outcome association (e.g., sex, month of birth) but we suspected many unmeasured confounders, *U*.

The UK Biobank study is a sample of 502, 644 UK residents enrolled between 2006 and 2010 [33]. At enrolment, the invited participants were aged between 40 and 69 years old and so would have turned age 15 between 1952 and 1985. The study response rate was 5.5% and higher levels of educational achievement predicted participation [34]. This suggests that the study participants were selected depending on *X*, educational attainment, which can bias an IV analysis (see earlier discussion and simulation study).

We performed 2SLS estimation using the linear probability model, where the exposure effect is on the risk difference scale [35]. Robust standard errors were calculated to account for assumptions about homogeneous exposure effects and the outcome distributions. For comparison, we also considered the equivalent standard analysis; that is, the linear regression of *Y* on *X* also with robust standard errors. Although a linear regression may be biased by unmeasured confounding of the *X* – *Y* association, we know from the missing data literature (e.g., [36]) that its exposure effect estimate is not biased by selection on *X*.

We used inverse probability weighting [37] to account for selection on educational achievement; thus the weighted IV analysis accounts for unmeasured confounding and nonrandom selection [20]. The weights were generated under the assumption that selection only depended on *X*. Those participants suspected to be under-represented in the selected sample (i.e., left school age 15) had larger weights, and hence contributed more to the weighted analysis, than those suspected to be over-represented in the selected sample (i.e. left school age 16 or older). For comparison, we carried out a weighted linear regression analysis using the same weights.

Table 2 presents the results for the exposure effect estimated using un-weighted and weighted versions of linear regression and IV analysis. For the IV analysis there were noticeable differences between the unweighted and weighted analyses. For outcome “ever smoker”, the weighted IV estimate was more than double that of the unweighted IV estimate, although there was some overlap between the corresponding 95% CIs. Both analyses suggested staying in school at least one extra year decreased the likelihood of being an ever smoker compared to those who left school at age 15, although the CI for the unweighted analysis was inconclusive since it included all 3 possible conclusions: risk decrease, no effect, and risk increase. For outcome “current smoker”, the results of the unweighted IV analysis suggested staying in school at least one extra year increased the likelihood of being a current smoker compared to those who left school at age 15, whilst the results of the weighted IV suggested the opposite effect. The CI for the unweighted analysis was inconclusive, including all 3 possible conclusions. As expected the unweighted and weighted linear regression results were identical.

**Table 2.** Risk difference %, of ever smoker or current smoker, for leaving school at age 16 or older compared to leaving school at age 15 using unweighted and weighted versions of linear regression (LR) and instrumental variable (IV) analysis. 95% confidence intervals displayed within brackets.

| Analysis | Outcomes |  |
| --- | --- | --- |
|  | Ever smoker | Current smoker |
| LR | -20.5 (-22.8, -18.3) | -14.1 (-15.5, -12.7) |
| Weighted LR | -20.5 (-22.8, -18.3) | -14.1 (-15.5, -12.7) |
| IV | -4.8 (-11.6, 1.9) | 1.8 (-1.5, 5.0) |
| Weighted IV | -10.6 (-14.8, -6.4) | -4.5 (-6.6, -2.4) |

Comparing the analyses which should not be biased by selection on *X*, the linear regression exposure effect estimates were about 2 to 3 times larger than those of the weighted IV, and there was no overlap in the 95% CIs. These differences may be due to the presence of unmeasured confounding which would only bias the linear regression analyses. However, other possible causes of the differences include an instrument that does not satisfy the IV analysis assumptions or heterogeneous treatment effects.

## Discussion

For nine different selection mechanisms, we have explained the structure of the selection bias and showed how DAGs can be used to determine if selection violates any of the IV assumptions. The IV estimate of the causal exposure effect is not biased by selection when selection is completely at random, depends only on the instrument, or depends only on unmeasured confounders. For the remaining selection mechanisms we have illustrated, using simulations, that nonrandom selection can result in a biased IV estimate and CI undercoverage. For a causal and null exposure effect, the IV estimate was biased, with often poor to severe CI undercoverage, when selection depended on the instrument plus measured confounder, or depended (in part or entirely) on the exposure, or depended on the outcome plus exposure. A special case was selection depending on the outcome only, where the IV estimate was only biased when *X* truly caused *Y*. Decreasing the instrument strength resulted in an increase in the level of bias for selection mechanisms partly depending on the instrument, but had little effect on the other selection mechanisms. For all selection mechanisms, CI coverages were noticeably higher for the moderate instrument compared to the strong instrument because standard errors increase with decreasing instrument strength. Whilst the larger standard errors improved CI coverage, there was still substantial CI undercoverage. Estimating the conditional IV estimate eliminated selection bias when caused by measured confounding, but only reduced the level of bias when selection resulted in measured and unmeasured confounding. Changing the exposure-instrument association from linear to nonlinear (i.e., cubic) reduced the size of the standard errors, but its effect on bias depended on the structure of the selection bias.

In keeping with the results of our simulation study, non-trivial levels of selection bias were demonstrated via simulations [24, 23, 22, 19, 21, 18]. [18] investigated two selection mechanisms in the context of Mendelian randomisation, and the remaining papers only considered a specific selection scenario.

Nonrandom selection can occur in practice, with large differences in the characteristics of the selected and unselected participants, as in our simulations. For example, the percentage of subjects who owned their property outright was 56.7% in the UK Biobank study (i.e., the selected sample) and 40.6% in the 2001 UK census (i.e., the study population) [34], so the odds of selection among outright property owners was almost double that of those who were not outright property owners. Similarly, using similar calculations for the Avon Longitudinal Study of Parents and Children (ALSPAC) study [38], the odds of selection among households with a car was almost double the odds of selection among households without a car.

Our simulation study has several limitations. First, whilst we considered eight plausible selection mechanisms, it was not possible to investigate all possible selection mechanisms even for a single IV analysis example. Second, in practice an IV analysis may use weaker instruments than we considered. We chose a sample size that was typical of an IV analysis so that even for an partial 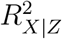 of 0.045 the instrument would not be considered weak. However, for the purposes of our study, we wanted to ensure that any bias was attributable only to selection and not to weak instrument bias [30]. Third, our simulation study was designed to show the effects of different selection mechanisms on an IV analysis and not an exhaustive investigation of the levels of selection bias that could occur in practice. Fourth, our use of non-parametric DAGs, to determine if selection would violate one of the core IV assumptions, are not suitable for all types of selection mechanisms (e.g., when the occurrence of selection bias depends on the parametrisation of the IV analysis [8]).

Some selection mechanisms bias the IV estimate but not the usual regression estimate; for example, when selection depends on the exposure. Unlike the IV estimate, the usual regression estimate may be biased by confounding-but the selection bias (for the IV estimate) may exceed the bias due to confounding of the usual regression estimate. Inverse probability weighting [21] and multiple imputation [23] have been used to appropriately account for selection in an IV analysis. Commercially available implementations of these methods usually assume the chance of selection depends on measured variables that are fully observed for all participants of the full sample. When selection depends on unmeasured variables or partially observed variables, MI and IPW may not fully account for selection and so give a biased IV estimate [23, 20]. There are some approaches to account for selection depending on unmeasured or partially observed data [23, 20, 22], but they tend to be less straightforward, make untestable assumptions or are specific to a particular type of IV analysis.

With individual-level data on the selected and unselected participants, an IV analyst can investigate possible factors that influence selection. However, this is impossible when the IV analyst only has summary-level data. Providers of summary-level data should discuss whether the study sample is a nonrandom sample of the target population, and posit possible selection mechanisms. Where possible, these providers could generate summary-level data accounting for nonrandom selection (e.g., summary-level data from a weighted analysis, or summary-level data adjusted for known factors associated with selection). Two-sample IV analyses tend to be conducted using summary-level data, and these analyses are further complicated because there are two opportunities for nonrandom selection to occur, and possibly two different selection mechanisms to take into account.

In summary, ignoring how participants are selected for analysis can result in a biased IV estimate, substantial CI undercoverage, and lead to an incorrect conclusion that an exposure is/is not causal. This limitation should be more widely noted in guidelines for IV analyses. DAGs can be used to assess if the IV analysis may be biased by the assumed selection mechanism. Future work could provide researchers guidance on statistical methods, diagnostic tools and sensitivity analyses for estimating causal effects from nonrandom samples.

